# Integration of single-cell transcriptome and genetic profiles reveals critical cell types and genes in inflammatory bowel disease

**DOI:** 10.64898/2025.12.06.692714

**Authors:** Xiaojing Chu, Siyu Wang, Jiacong Wei, Haosheng Liu, Pengcheng Yan, Qiuli Yang, Yu Tang, Bowen Zhang, Erli Pang

**Author notes:** These authors contributed equally (Xiaojing Chu, Siyu Wang, Jiacong Wei). Correspondence (X.C.), (E.P.).

## Abstract

The progression of inflammatory bowel disease (IBD), grouped into Crohn’s disease (CD) and ulcerative colitis (UC), involves multiple cell types and genes, but the identification of critical cell types and genes that contribute to intestinal inflammation remains challenging. By integrating human intestinal single-cell transcriptomic data from approximately 200 donors, we revealed the stromal cell-derived network in CD, enrichments of immune cells such as Th1 in CD, atypical B cells and Tfh in UC*, AXL*^+^*SIGLEC6*^+^ DCs and IgG+ Plasma B cells in both, and the association between *SPP1*^+^ Macrophages, *FAP*^+^ Fibroblasts and anti-TNF treatment resistance. We also investigated the gene expression characterization of enteric neurons. Additionally, by linking cell-type–specific transcriptional alterations with genetic profiles of disease risk and intestinal gene expression, we proposed B cells and cycling epithelial cells as critical cell types, and identified epithelial genes, such as *GPX4,* as associated with CD pathology. In summary, our study comprehensively characterized the single-cell landscape of human intestines in IBD and provided insights into cell types and genes highly involved in genetically determined CD pathology, thereby informing CD prediction, diagnosis, and the development of new therapeutic targets.

## Introduction

Inflammatory bowel disease (IBD) is the most common intestinal immune-mediated disease grouped into Crohn’s disease (CD) and ulcerative colitis (UC). Among them, CD affects both large and small intestines and exhibited higher genetic determinants^1^. UC is limited to the colon, same to colorectal cancer (CRC). Large-scale genome-wide association analyses have identified hundreds genetic risk loci^2–4^, and the enrichments of inflammatory regulators and epithelial function genes highlight the pathological roles of different human intestinal components. Meanwhile, gene expression profiles at the single-cell level have also reported changes in multiple cell-type compositions, covering epithelial, stromal, and immune cells, and uncovered gene programs showing alterations in inflammatory status^5,6^ and relevance to clinical treatment^7,8^.

However, the previous studies have largely focused on either genetic profiles to map the cell types and genes directly, thus lacking the evaluation of their effective roles in active contexts, or transcriptional alterations, which may not reflect causality in strongly genetically driven diseases. Therefore, it remains a challenge in linking genetic regulation profiles with single-cell transcriptomic profiles to illustrate cellular and molecular mechanisms critical to IBD onset and development in an integrative study.

Integrating large-scale single-cell transcriptomic data greatly enhanced the power in identifying rare cell subsets, and diminished the discrepancy introduced by different analysis procedures from different studies, resulting in a more comprehensive and generalized portrait. Additionally, leveraging the cellular characterization from single-cell transcriptomics, an opportunity was acquired to redefine cellular and molecular involvements for genetic profiles and has achieved remarkable results in limited attempts^9^.

In this study, by first integrating single-cell data from multiple datasets, we constructed a harmonized intestinal cell atlas covering samples of different intestinal regions from healthy volunteers, patients with CD and UC. We systematically compared the cellular composition of large and small intestines under different pathological states and characterized the cellular relationships with stromal cells as network hubs that may lead to dysregulation in CD. We revealed the immune cell functional remodeling in broad human intestinal diseases, presenting a full landscape of human intestinal immunity in both hyper-response and suppression. We showed the power of our established atlas in characterizing rare cell types for broad applications. Furthermore, we prioritized cell types and genes critical to CD by integrating cell-type-specific transcriptional alterations with genetic profiles of disease risk and intestinal gene expression. In summary, these findings provide valuable insights into IBD pathology from disease associated cell networks, cell compositions and gene expression alterations, and genetic regulation, which may guide the development of preventive medicine, biomarkers, and new therapeutic targets.

## Results

### A single-cell atlas of inflammatory bowel disease

To establish a comprehensive single-cell atlas of IBD, we collected published scRNA-seq data covering healthy intestinal tissues, uninflamed and inflamed tissues of CD and UC^5,7,8,10–15^, containing 442 samples from either large intestines or small intestines of 196 donors with or without treatment (Fig. 1a,b, Supplementary Table 1). Among the 120 patients, who suffered from CD, 73 patients were reported to be under immune-inhibitor treatment. After quality control largely consistent with our previous study on human colorectum^16^, we classified 713,612 cells of high-quality into 71 subsets, including 12 epithelial cell subsets, six endothelial cell subsets, nine fibroblast or pericyte subsets, two neuronal cell subsets, seven B cell subsets, 16 T cell subsets, four NK/ILC subsets, and 15 myeloid cell subsets, based on their distinct expression patterns (Fig. 1c, Extended Data Fig. 1-3). To address the concerns of batch effects, we showed that the cells were clustered based on cell types rather than datasets (Extended Data Fig. 4a). By further performing a principal component analysis (PCA) on all samples based on their cell subtype proportion in either all non-epithelial cells or all immune cells, the results supported that samples of different datasets, but the same pathological states and sampling regions were clustered together (Extended Data Fig. 4b-d). We also observed that B and T cell subsets contributed the most variation (Supplementary Table 2), in line with their pathological roles in IBD. For the application in a broader field, we established an online browser https://cmb.bnu.edu.cn/Chulab_CD.

**Fig. 1.**
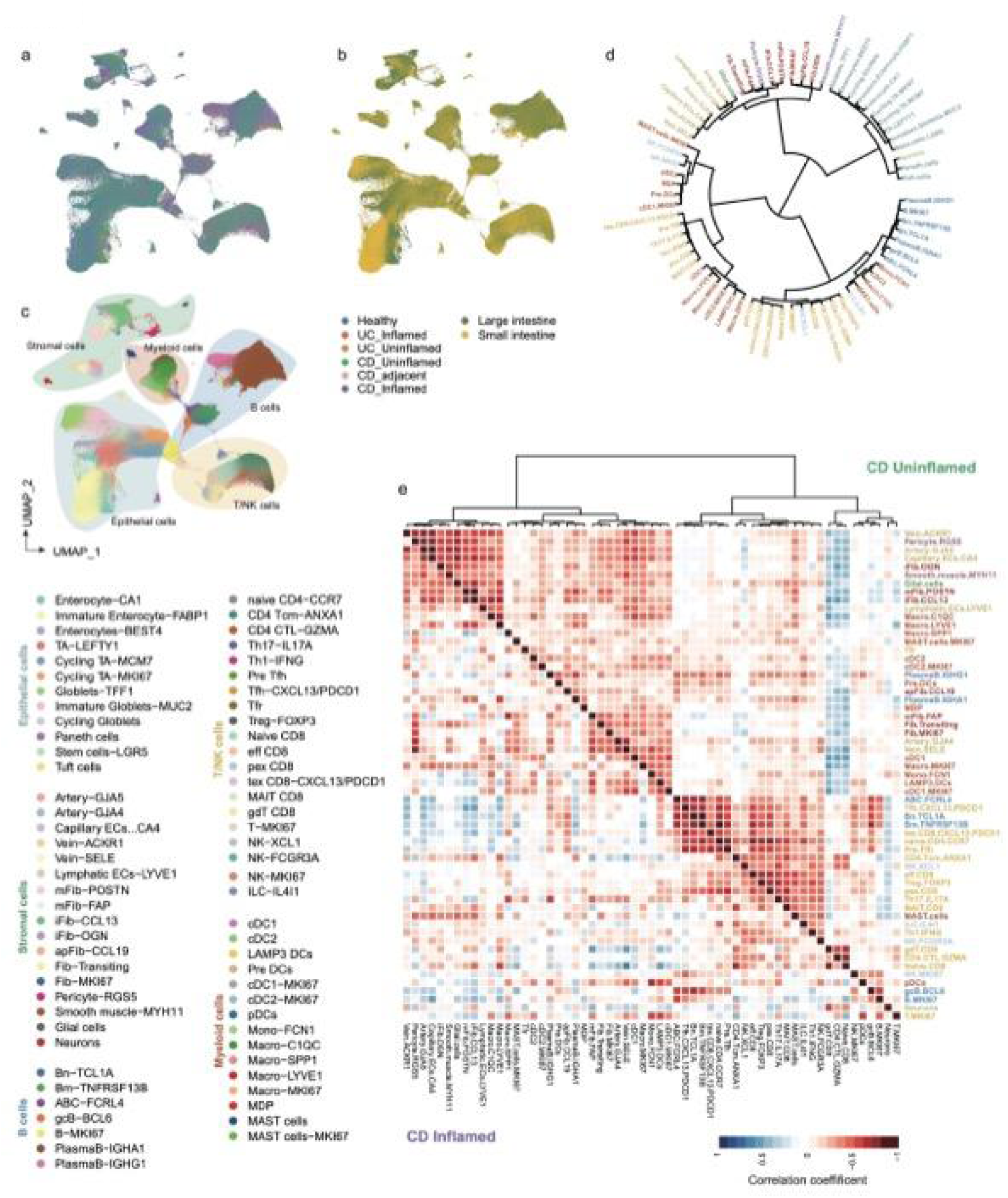
Single cell landscape of human intestines. **a-c**, UMAP plot showing cell clustering colored by tissue pathological states (a), sampling regions (b) and identified subsets (c) (n=713612 cells). fib: fibroblasts, n: naïve, m: memory, cm: central memory, gc: germinal center, macro: macrophage, eff: effector, pex: progenitor exhausted, tex: terminal exhausted. **d**, Hierarchical clustering of cell subsets based on average expression levels in all samples (n = 314 samples). **e**, Hierarchical clustering of cell subsets based on *Spearman’s* correlation coefficients between each of two subsets in either CD uninflamed samples (top right, n = 124 samples) or CD inflamed samples (bottom left, n = 68 samples).

Inspired by some cell subsets of different cell lineages clustered together at the transcriptomic level (Fig. 1d), we next investigated the cellular relationships by clustering cell subsets based on their composition in different samples. Consistent with their cellular origins, cell subpopulations from the same lineage tended to be clustered together in all samples (Extended Data Fig. 5a). However, different from the stronger cell occurrence observed in tumor samples reported previously^16^, in contrast to the uninflamed samples from CD patients, the cellular occurrence was markedly reduced in inflamed tissues (Fig. 1e). For example, the negative correlations between stromal cells and T cell subsets were largely missing, suggesting a dysregulation in CD.

Further, to infer the potential cause of the dysregulation, we performed a cell communication analysis in each pathological status and more specific interactions were detected in inflamed samples of CD patients (Fig. 2a) than in uninflamed samples of IBD patients, inflamed samples of UC patients as well as samples from healthy donors (Extended Data Fig. 5b), suggesting that a hyper-activated cell network may be associated with the dysregulation in CD. In addition, we identified stromal cells, such as pericytes and fibroblasts, as the major sending cell types with the most ligands expressed, and immune cell types as the major targets in the communications (Fig. 2b, Extended Data Fig. 5c). We also observed that fibroblasts and perivascular cells in the inflamed samples of CD upregulated gene pathways related to stromal cell development and response to TNF signaling (Fig. 2c, Extended Data Fig. 5d). Particularly, LAMB3-mediated cell signaling was identified in the perivascular cells of inflamed samples of CD (Fig. 2d), where *LAMB3* was exclusively expressed (Fig. 2e), in line with the reported LAMB3 effects in promoting intestinal inflammation^17^. We next validated the predicted interactions between *LAMB3* and *CD44* in spatial transcriptomic data of inflamed tissues of CD^6^ with a significant colocalization (*Fisher’s* exact test, p = 3.20e-13, Fig. 2f and Extended Data Fig. 5e). Together these results revealed a stromal cell-derived cell network in CD, which may lead to the loss of cell occurrence.

**Fig. 2.**
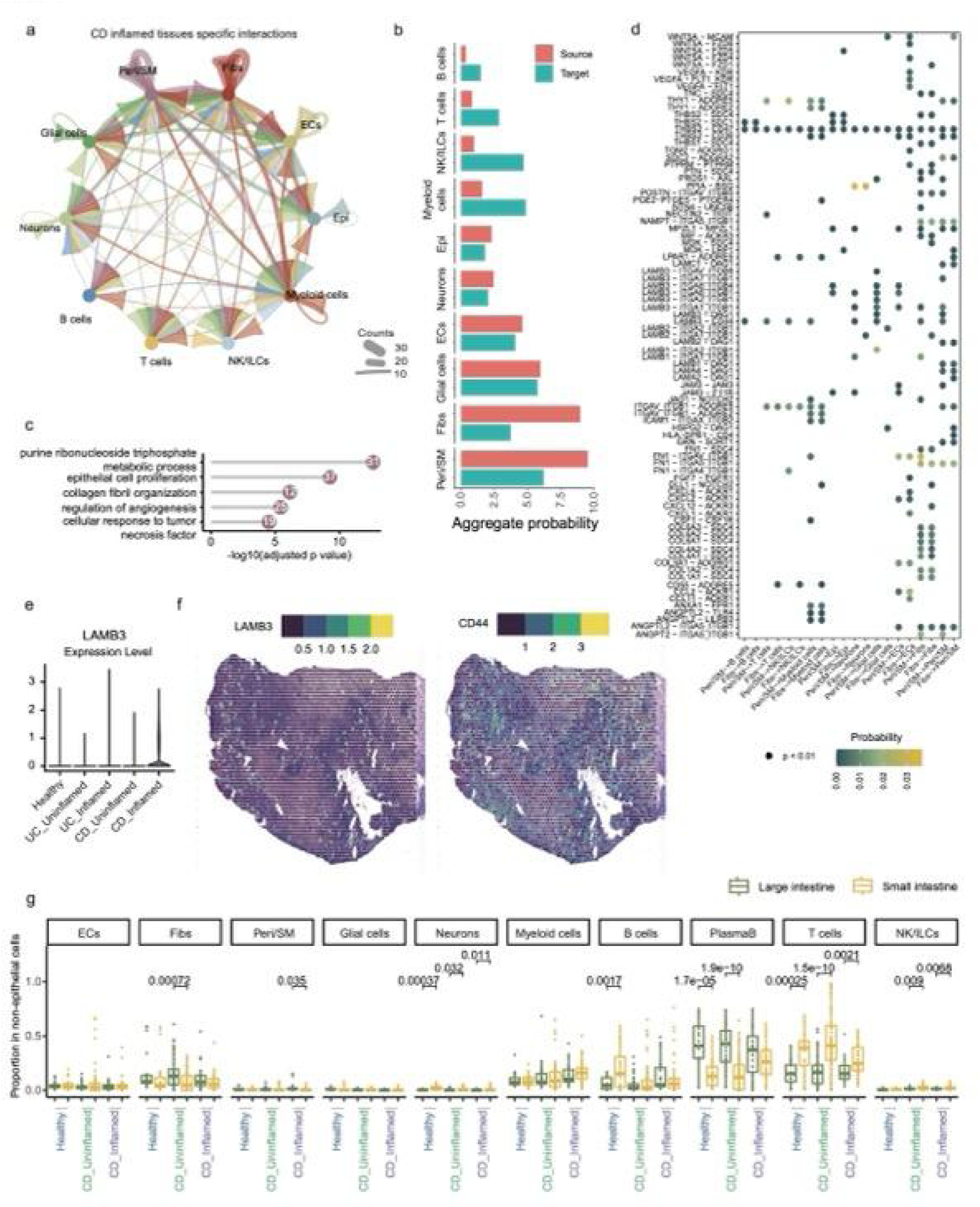
Inferred cell interactions in CD inflamed tissues. **a**, Circle plot showing counts of predicted cell interactions specific to CD inflamed samples. **b**, Barplot showing aggregate probabilities of each cell type as source and target in cell interactions. **c**, Lollipop plot showing upregulated pathways in the inflamed tissues compared to uninflamed tissues in pericytes. Numbers in the dots indicate gene counts matched to corresponding biological pathways. Over-representation analysis (BH adjustment). **d**, Dotplot showing estimated probability of ligand-receptor pairs from fibroblasts and pericytes to other cell types identified in only CD inflamed tissues. **e**, Violin plot showing the expression level of *LAMB3* in pericytes of different tissues (n = 658 (Healthy), 138 (UC_Uninflamed), 278 (UC_Inflamed), 1412 (CD_Uninflamed) and 2036 (CD_Inflamed) cells). **f**, Colocalization of *LAMB3* and *CD44* in inflamed regions revealed by spatial transcriptomics. **g**, Boxplots showing cell type proportion in all non-epithelial cells (n = 21, 26, 40, 84, 27 and 41 from left to right). Center line indicates the median value, lower and upper hinges represent the first and third quartiles, respectively, and whiskers denote 1.5× interquartile range. *Student’s* t-test (two-sided, unpaired).

### Myeloid cells remodeling in intestinal diseases

We next compared the major cell type preference between samples of the large intestine and the small intestine. As shown in Fig. 2g, significant differences were observed in the cell proportion of neurons, T cells and NK/ILCs with larger abundances in small intestine and plasma B cells with larger abundances in the larger intestine. Additionally, the proportion of plasma B cells was decreased in the inflamed regions compared to controls in large intestine, while increased in small intestine.

Considering the significant influence of sampling regions, we identified cell subsets associated with IBD in only large intestine samples through comparisons conducted among different pathological states. We observed distinct cell composition alterations (Extended Data Fig. 6a, Supplementary Table 3). As presented in Fig. 3a, myeloid cells exhibited a higher proportion in CD but a lower proportion in UC compared to healthy colorectal tissues. In line with the previous result about stromal cell-derived network, we also observed an active cell-cell interactions between myeloid cells and stromal cells (Fig. 3b). Notably, myeloid cells in the inflamed samples of CD patients highly expressed *CXCL8/9* and *CCR1* (Fig. 3c), contributing to an activation of cell communications with stromal cells specific to CD (Fig. 3d). It was reported that CCR1 mediated the accumulation of myeloid cells in tumors^18,19^, and the role of CXCL8/9 in enhancing stromal cell proliferation^20^. Together with our observation that myeloid cells in inflamed regions upregulated pathways related to lymphocyte activation (Fig. 3e), we highlighted the pathological roles of myeloid cells in contributing to CD pathology.

**Fig. 3.**
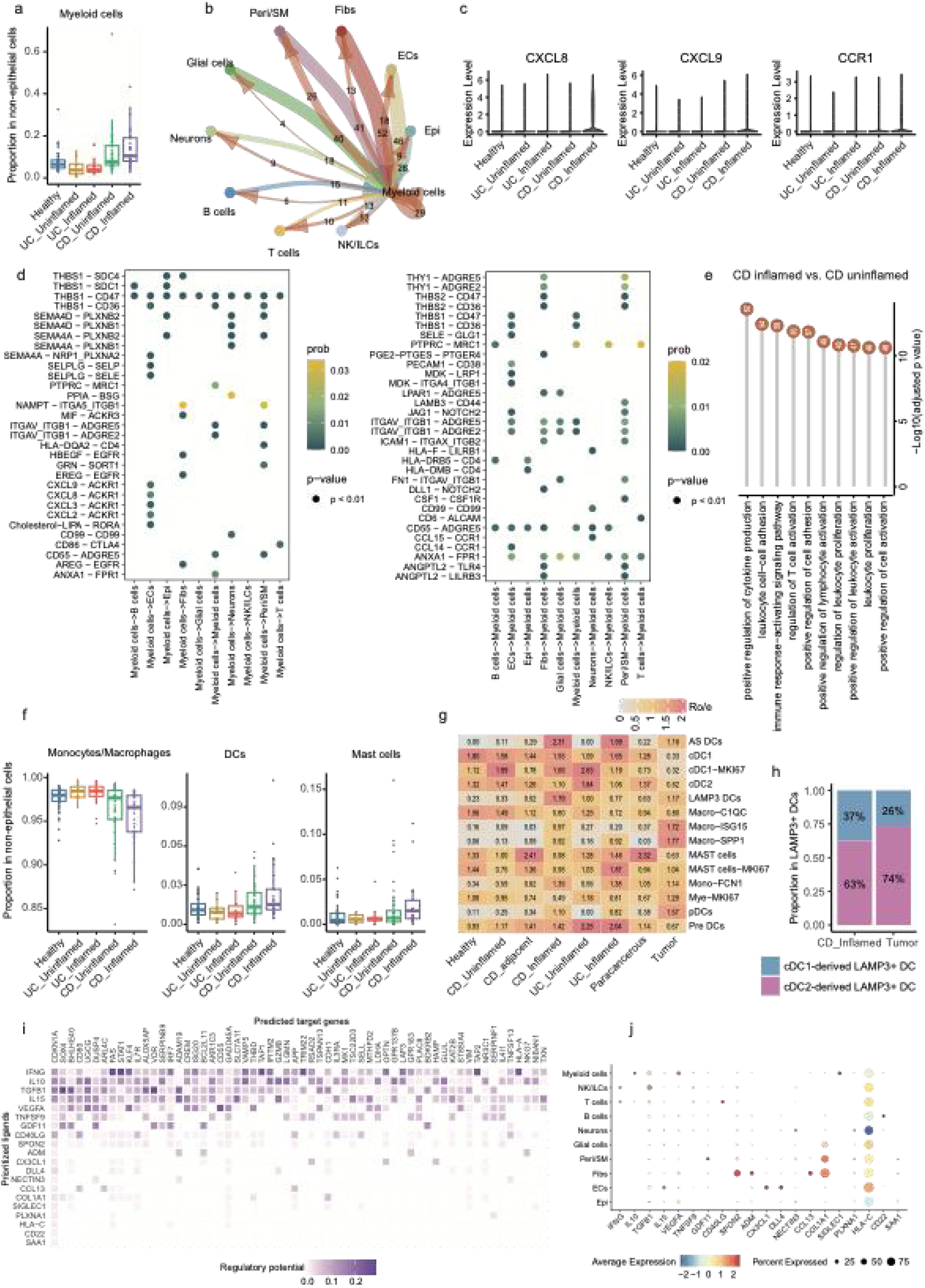
Myeloid cells characterization in human intestinal diseases. **a**, Boxplot showing large intestinal myeloid cell proportion in all non-epithelial cells (n = 45, 14, 18, 40 and 27 samples from left to right). Center line indicates the median value, lower and upper hinges represent the first and third quartiles, respectively, and whiskers denote 1.5× interquartile range. *Student’s* t-test (two-sided, unpaired). p values were presented in Supplementary Table 3. **b**, Circle plot showing counts of all predicted interactions around myeloid cells. **c**, Violin plots showing the expression level of *CXCL8/9* and CCR1 in myeloid cells from different tissues (n = 6297, 797, 882, 19353 and 18620 cells from left to right). **d**, Dotplots showing estimated probability of ligand-receptor pairs from (Left) or to (Right) myeloid cells identified in only CD inflamed tissues. **e**, Lollipop plot showing upregulated pathways in the inflamed tissues compared to uninflamed tissues in myeloid cells. Numbers in the dots indicate gene counts matched to corresponding biological pathways. Over-representation analysis (BH adjustment). **f**, Boxplots showing myeloid cell subpopulations proportion in all non-epithelial cells in large intestine samples (n = 45, 14, 18, 40 and 27 from left to right). Center line indicates the median value, lower and upper hinges represent the first and third quartiles, respectively, and whiskers denote 1.5× interquartile range. *Student’s* t-test (two-sided, unpaired). p values were presented in Supplementary Table 4. **g**, Heatmap showing tissue preference of each myeloid cell subset estimated by *Ro/e* score. **h**, Stacked bar plots showing the predicted origins of LAMP3^+^ DC in different tissue types. **i**, Heatmap showing the regulatory potentials of the top 20 prioritized ligands of AS DCs. **j**, Dotplot showing the expression levels of the top 20 prioritized ligands in each cell type.

By next zooming in on myeloid cell subpopulations, we found that DCs and mast cells drove the observation in all non-epithelial cells, showing enrichments in CD (Fig. 3f, Supplementary Table 4), in line with the previous report^5^. The potential important roles of DCs indicated by the result encouraged us to further investigate the functional preference of myeloid cells in a broad human intestinal disease context. To cope with it, we integrated the IBD atlas with colorectal cancer (CRC) data extracted from our previous established single-cell atlas of human colorectum^16^ (Extended Data Fig. 6b). By showing the tissue preference of the myeloid functional subsets well-recognized by our and other previous studies^21,22^, we found LAMP3^+^ DCs were enriched in cancers as well as in CD inflamed tissues (Fig. 3g), supporting the previous reports^23–27^. Studies in tumor microenvironments indicated multiple cell origins of LAMP3^+^ DCs, corresponding to their diverse function potentials^28^. By aligning LAMP3^+^ DCs to their two reported major cellular origins, cDC1 and cDC2, we found a higher proportion of cDC1-derived LAMP3^+^ DCs in CD than in cancers (Fig. 3h), agreeing with the more immune suppressive roles of cDC2-derived LAMP3^+^ DCs^28^. We also observed more activation of antigen processing functions in cDC1 of CD than in tumors (Extended Data Fig. 6c), which may explain the enrichment of cDC1-derived LAMP3^+^ DCs in CD.

Additionally, we found that AS DCs, marked by the exclusive expression of *AXL* and *CD5* and the high expression of *CD80, SPI1, KLF4* and *RUNX3* (Extended Data Fig. 6d), were highly enriched in IBDs than in cancers (Fig. 3g), in line with their reported pro-inflammatory roles^22^. To explore the reason of AS DCs enrichment in inflammation context and consider AS DC was reported as a transiting state between cDC2 and pDC^21^, extracellular regulators were predicted by NicheNet analysis^29^ on target genes identified using cDC2 and pDCs as controls. We found, for examples, *IFNG*, *IL10* mainly expressed by T/NK and myeloid cells, and *CCL13* mainly expressed by fibroblasts may be responsible for the unique phenotype of AS DCs in CD (Fig. 3i,j and Extended Data Fig. 6e). Among them, fibroblast-*CCL13* has been linked to DCs with pro-inflammation roles in autoimmune diseases^30,31^.

### Lymphocyte subsets preference in intestinal diseases

Next, we examined the tissue distribution preference of each identified cell subset with Ro/e analysis^32^ and observed more universal alterations in the large intestines than that in the small intestines (Fig. 4a, Extended Data Fig. 6f). Notably, different from that in CRC, where stromal cells exhibited large-scale alterations^16^, most changes were observed for immune cell components in IBD (Fig. 4a,b), consistent with their dominant contribution to PCA (Supplementary Table 2), indicating the effective roles of immune cells in intestinal inflammation.

**Fig. 4.**
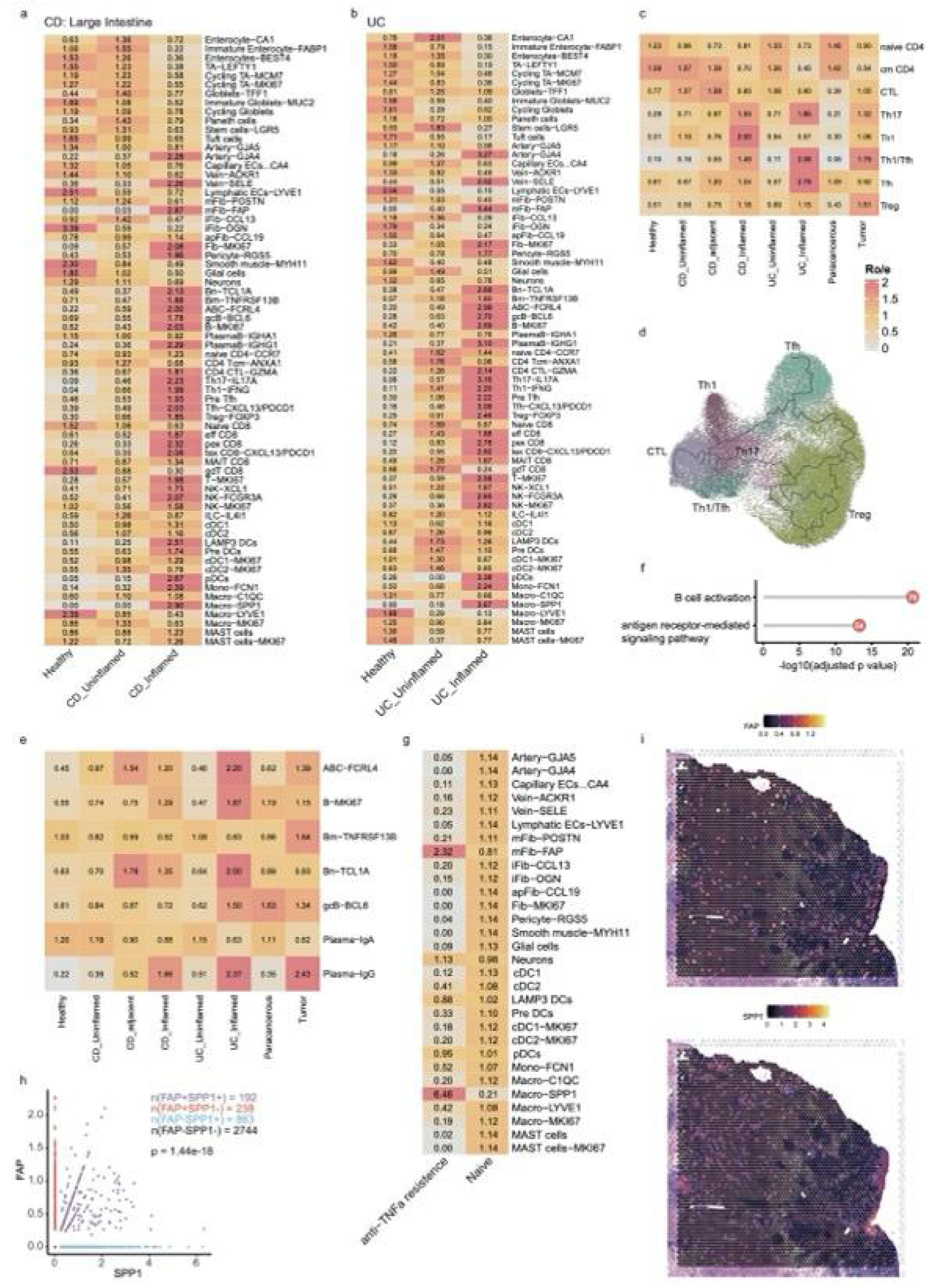
Lymphocyte subsets preference in human intestinal diseases. **a-c**, Heatmap showing tissue preference of each lymphocyte subset estimated by *Ro/e* score. **d**, Predicted cell trajectory of activated CD4^+^ T cells (n = 67532 cells). **e**, Heatmap showing tissue preference of each B cell subset estimated by *Ro/e* score. **f**, Lollipop plot showing upregulated pathways in ABCs compared to other B cell subsets. Numbers in the dots indicate gene counts matched to corresponding biological pathways. Over-representation analysis (BH adjustment). **g**, Heatmap showing tissue preference of each cell subset estimated by *Ro/e* score. **h**, Scatter plot showing expression of *FAP* and *SPP1* in spatial spots of all slides. *Fisher’s* exact test. **i**, Colocalization of *FAP* and *SPP1* in inflamed regions revealed by spatial transcriptomics.

It was reported that CD4^+^ T cells were pathologically involved in both IBD and CRC as key regulators of immune tolerance and inflammation^33–37^. Therefore, we integrated all CD4^+^ T cells from IBD atlas with CD4^+^ T cells of paracancerous tissues and tumors from CRC atlas^16^ (Extended Data Fig. 7a). As expected, Th1 marked with the expression of *IFNG* was highly enriched in autoimmune (CD), and Treg was enriched in cancer, contributing to an immunosuppressive microenvironment (Fig. 4c). Inspired by the chronic intestinal inflammation may develop into CRC, we hypothesized a transition from other functional CD4^+^ T cell subsets to Treg, which was supported by the trajectory analysis showing the continuous embedding from Th17 and Tfh to Treg (Fig. 4d). The inferred transiting relationships agree with the reports that Th17 transdifferentiated into Tregs in a mouse autoimmune disease model^38^ and Treg transformed into Tfh in a mouse skin inflection model^39^.

B-cell components were also largely reprogrammed in both UC and CD inflamed tissues and had crucial functions in intestinal homeostasis^40–42^ and tumors^43,44^. Therefore, we redefined the tissue preference in the integrative IBD and CRC data (Extended Data Fig. 7b). We found that atypical B cells (ABCs) and IgG+ Plasma B cells were highly enriched in IBD inflamed regions and tumor tissues in contrast to normal tissues (Fig. 4e). Among them, ABCs upregulated pathways related to B cell activation and antigen signaling (Fig. 4f), agreeing with their proinflammatory roles^45^.

The data of anti-TNFa treatment response provided us an opportunity to investigate the cell subset related to immune inhibitor therapy resistance. By comparing with treatment naïve controls, non-responders highly enriched myofibroblast subset mFib-FAP and the macrophage subset Macro-SPP1 (Fig. 4g, Extended Data Fig. 7c). Macro-SPP1 was first identified as a subset of tumor-associated macrophages^46^ with functional potentials in angiogenesis and correlated with worse prognosis and immune-checkpoint blockade treatment resistance^47^. Inflammatory fibroblasts have been linked with anti-TNF resistance^6^. Supporting their potential interactions reported in CRC^48^, we observed the colocalization of spatial spots with *FAP* and *SPP1* signatures in CD inflamed tissue (Fig. 4h,i). To sum up, we revealed the remodeling of lymphocyte subsets in human intestinal diseases, may contributing to IBD and anti-TNF treatment resistance.

### Characterization of intestinal neurons in inflammation

The crucial roles of neurons in intestinal immune regulations have attracted attention^49^, however with few investigations in humans due to the sampling difficulties. Leveraging the single-cell integration technique, we revealed a cluster of intestinal neurons despite largely lost in IBD (Fig. 5a) and may not be recovered after immune inhibitor treatments (Extended Data Fig. 7d). A pathway enrichment analysis on upregulated genes in neurons from inflamed samples implied an enhanced oxidative phosphorylation and immune activation (Fig. 5b), marked by the high expression of class I MHC molecules (Fig. 5c), suggesting their pro-inflammatory roles. Next, we explored the intracellular regulations under these gene expression phenotypes, and uncovered transcription factors such as *STAT1, IRF9* that were activated and upregulated in intestinal inflammation (Fig. 5d, Extended Data Fig. 7e).

**Fig. 5.**
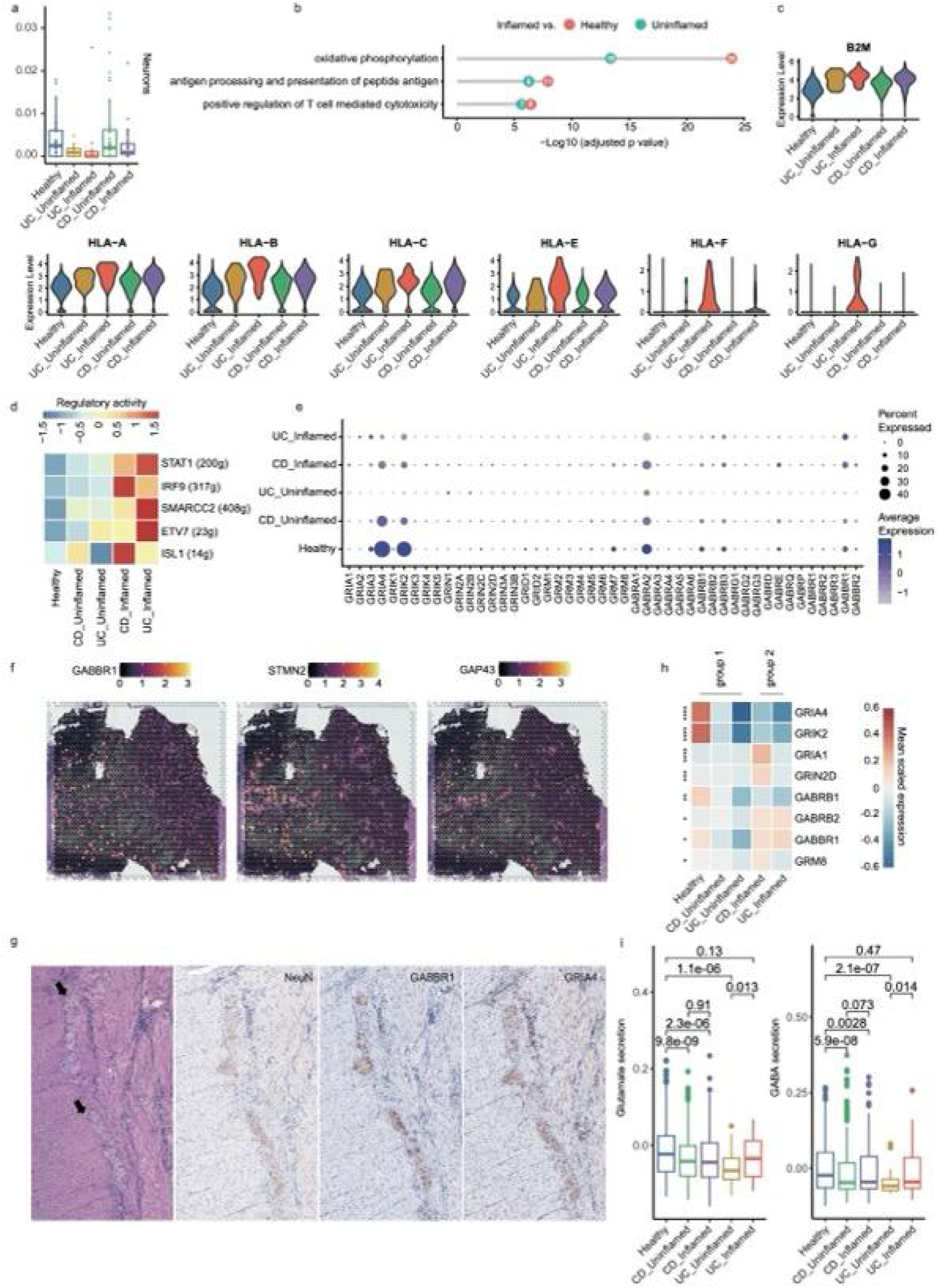
Intestinal neurons characterization. **a**, Boxplot showing large intestinal neurons proportion in all non-epithelial cells (n = 45, 14, 18, 40 and 27 samples from left to right). Center line indicates the median value, lower and upper hinges represent the first and third quartiles, respectively, and whiskers denote 1.5× interquartile range. *Student’s* t-test (two-sided, unpaired). p values were presented in Supplementary Table 3. **b**, Lollipop plot showing upregulated pathways in neurons of inflamed tissues compared to that of uninflamed and healthy tissues. Numbers in the dots indicate gene counts matched to corresponding biological pathways. Over-representation analysis (BH adjustment). **c**, Violin plots showing the expression level of MHC-I genes in neurons of different tissues (n = 714, 32, 37, 986 and 527 cells from left to right). **d**, Heatmap showing regulatory activity of representative transcription factors upregulated in inflamed samples. **e**, Dotplot showing expression levels of neuronal receptors in different tissues. **f**, Colocalization of *GABBR1* and reported top intestinal neuron markers revealed by spatial transcriptomics. **g,** Representative images (200x) of colorectal ganglion cells (indicated by arrows) stained by hematoxylin-eosin (H&E) and IHC. **h**, Heatmap showing expression levels of neuronal receptors differentially expressed in inflamed tissues compared to normal tissues. p values from top to bottom equal to 2.61e-13 (****), 2.67e-9 (****), 4.48e-7 (****), 9.09e-4 (***), 9.13e-3 (**), 3.07e-2 (*), 3.16e-2 (*) and 4.09e-2 (*). **i**, Boxplots showing signature scores of neurons from different tissues (n = 714, 32, 37, 986 and 527 cells from left to right). Center line indicates the median value, lower and upper hinges represent the first and third quartiles, respectively, and whiskers denote 1.5× interquartile range. *Student’s* t-test (two-sided, unpaired).

Considering the role of neurons as a regulation hub, we examined the expression of all glutamate and GABA receptor genes (Fig. 5e). We observed that glutamate AMPA receptor gene *GRIA4* and glutamate kainate receptor gene *GRIK2,* among all other ionotropic glutamate receptor genes, were highly expressed in human intestines, and several GABA receptor genes were also showed remarkable expression levels, in line with their alleviating inflammation in intestines^50^. By revisiting the spatial transcriptomic data of human intestine^6^, we validated the expression of *GRIK2*, *GABRA2*, *GABRB1*, *GABRB3*, *GABRE* and *GABBR1* in all spots with neuron signatures (Extended Data Fig. 7f) and showed the colocalization of *GABBR1* with reported top neuron markers in a representative slide as an example (Fig. 5f, Extended Data Fig. 7g). Immunohistochemistry (IHC) staining in human colon tissue samples further validated the expression of two representative neuronal transmitter receptors: GRIA4 and GABBR1 at the protein level (Fig. 5g). Notably, compared to neurons in uninflamed regions, *GRIA4*, *GRIK2* and *GABRB1* were downregulated and *GRIA1*, *GRIN2D*, *GABRB2*, *GABBR1* and *GRM8* were upregulated in inflamed regions (Fig. 5h), implying their involvement in neuron-immune interactions. The detection on gene signatures related to glutamate and GABA synthesis and secretion mirrored altered glutamate and GABA release in inflammation (Fig. 5i, Extended Data Fig. 7h). To sum up, our integrative single-cell atlas offered an opportunity to characterize rare cell type in human intestine.

### B cells and epithelial cells as CD critical cell types

Considering CD as a genetics-driven disease, we next aimed to identify cell types crucial to CD. By integrating GWAS profiles and single-cell profiles, we first applied two frameworks, as illustrated in Fig. 6a,b (also see Methods). a: (1) Cell type–specific genes and genes differentially expressed in CD compared to healthy controls (Disease-associated genes) were identified. (2) An enrichment analysis was performed on GWAS profiles as indicated by explained heritability conditional on single nucleotide polymorphism (SNP)s physically located in cell type–specific genes or disease-associated genes. b: (1) We mapped CD GWAS SNPs to genes according to different criteria. (2) Significant overlaps with cell type specific and disease associated genes were detected. Additionally, accentuating disease-associated transcriptional alteration, we also applied an approach, which was adjusted based on our previous study^16^ (Methods). Simplified as c: transcriptional alteration levels, defined by distribution deviation in differential expression analysis, were also estimated for all expressed genes and other predefined gene lists.

**Fig. 6.**
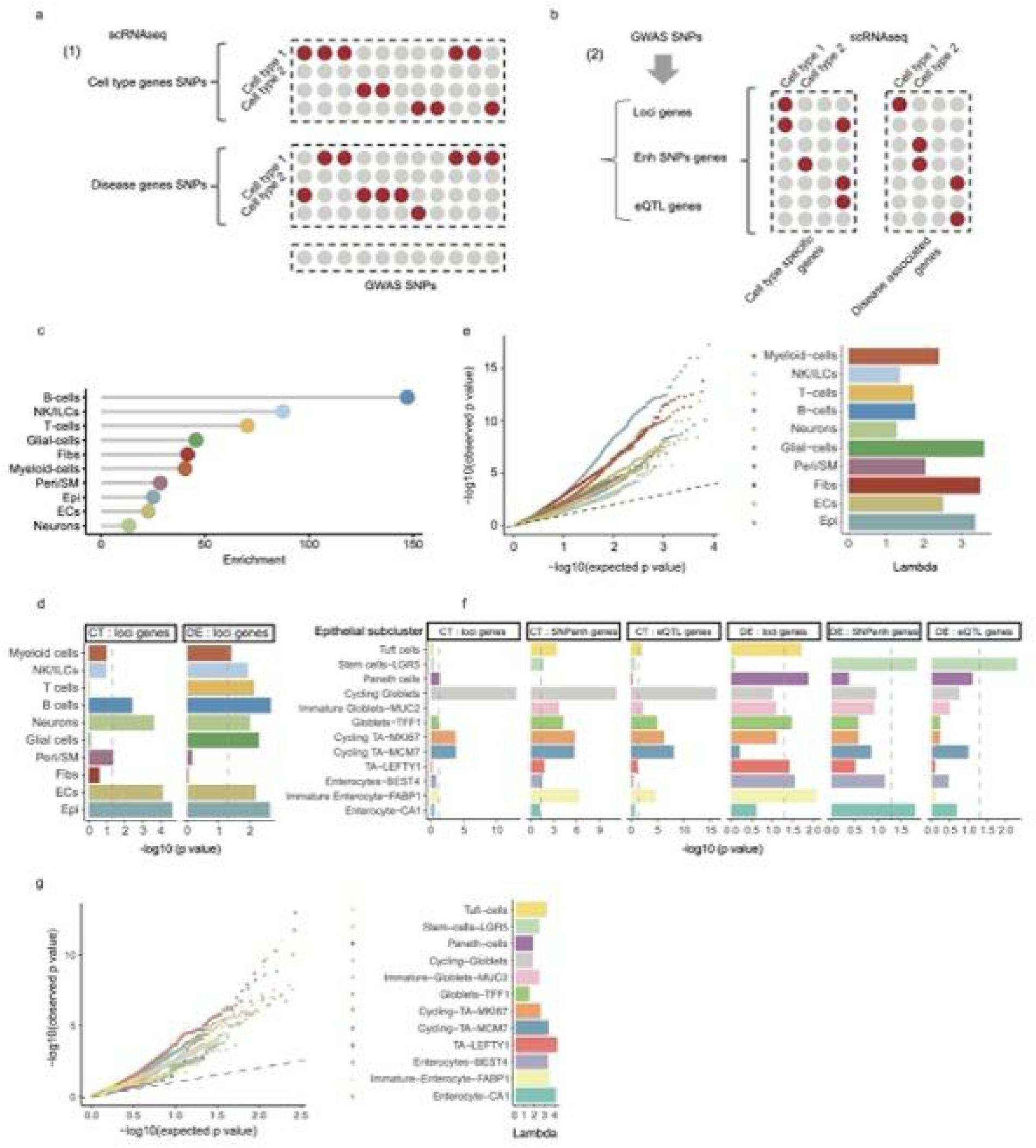
Identification of CD critical cell types. **a-b**, Schematics of CD critical cell type identification frameworks. **c**, Lollipop plot showing enrichment score in sLDSC analysis. **d**, Barplots showing enrichment significances of loci genes in cell type specific (CT) and disease associated (DE) genes. *Fisher’s* exact tests. **e**, Left: qqplot showing p value distribution of all expressed genes in the comparisons between CD inflamed tissues and healthy tissues (n = 8535, 2132, 7046, 10055, 2507, 1170, 1679, 8965, 7881 and 11838 genes from top to bottom). Right: Barplot showing lambda values of different cell types. **f**, Barplots showing enrichment significances of loci genes, genes with SNPs located in enhancer regions and eQTL genes in cell type specific (CT) and disease associated (DE) genes. *Fisher’s* exact tests. **g**, Left: qqplot showing p value distribution of expressed loci genes in the comparisons between CD inflamed tissues and healthy tissues (n = 45, 342, 31, 183, 375, 315, 380, 406, 396, 195, 281 and 278 genes from top to bottom). Right: Barplot showing lambda values of different epithelial subtypes.

Consistent with the report on UC^9^, B cells were prioritized as the top associated cell types in CD (Fig. 6c) using cell type specific genes in the conditional heritability analysis, however inconsistent results were acquired with disease-associated genes (Extended Data Fig. 8a), supporting NK/ ILCs as critical cell types associated with the genetic risk of CD. Following the steps of framework b, we first identified epithelial cells and B cells as the top effector cell types with the most significant enrichment of GWAS loci genes (Fig. 6d). Then the distribution analysis in both all expressed genes and GWAS loci genes consistently supported the large alteration in CD epithelial cells (Fig. 6e, Extended Data Fig. 8b). Next, by zooming into epithelial cell subsets, framework b was applied on the combinations of cell type–specific genes and disease-associated genes with loci genes and genes covering either intestinal expression quantitative trait loci (eQTL) or SNPs in enhancer regions identified in human intestinal samples (Fig. 6f). Overlaps with cell type specific genes consistently supported a high relevance of genetic risk genes to cycling epithelial cells. Analysis on disease-associated analysis (Fig. 6f), together with distribution analysis on differential expression analysis profiles with both GWAS loci genes (Fig. 6g) and eQTL genes (Extended Data Fig. 8c) proposed TA, intestinal stem cells and enterocytes exhibited enriched transcriptional alterations in CD. Summarizing from all evidence, we proposed B cells and epithelial cells as critical cell types in CD.

### Identification of CD critical genes

Prioritizing CD-associated genes is an important task for both understanding CD pathological mechanisms and finding therapeutic targets. We followed two routes starting from either GWAS genes or cell type and disease genes (Fig. 7a,b). By intersecting GWAS loci genes with both cell type–specific and disease-associated genes, we acquired 185 unique genes for different cell types. Considering the previous clues of CD critical cell types, we focused on B cells and epithelial cells in the further filtering. Leveraging the information of membrane protein, eQTLs and enhancer regions, we proposed five B cell genes and two epithelial genes with top priority in CD pathology (Fig. 7a, Supplementary Table 5). Among them, *ICAM3* has been associated with diffuse large B-Cell lymphoma^51^, *CCR10* expressed on plasma cells was involved in CRC^52^, colonic homeostatic regulation^53^, B cell epithelial cell interactions^54^ and virus infection^55^. *GPX4* was reported to regulate mucosal epithelial cell ferroptosis^56,57^, supporting their potential inflammatory roles in CD. Lymphotoxin beta receptor (*LTBR*), linked to important intestinal epithelial immune functions^58^, has also been prioritized. Further, supporting their associations between genetic profiles of CD and *GPX4, LTBR* expression in ileum tissues, we observed strong colocalizations in the regions around rs4806969 (*GPX4*) and rs7954567 (*LTBR*), with H4 = 0.792 and H3+H4 = 0.958, respectively (Fig. 7c). Additionally, considering the high relevance of metabolic pathways to CD and genetic factors regulating metabolites levels enriched for nonsynonymous variants^59^, we also provided a gene set covering either variants of nonsynonymous SNV, stop-gain, stop-loss or frameshift substitutions (Supplementary Table 5).

**Fig. 7.**
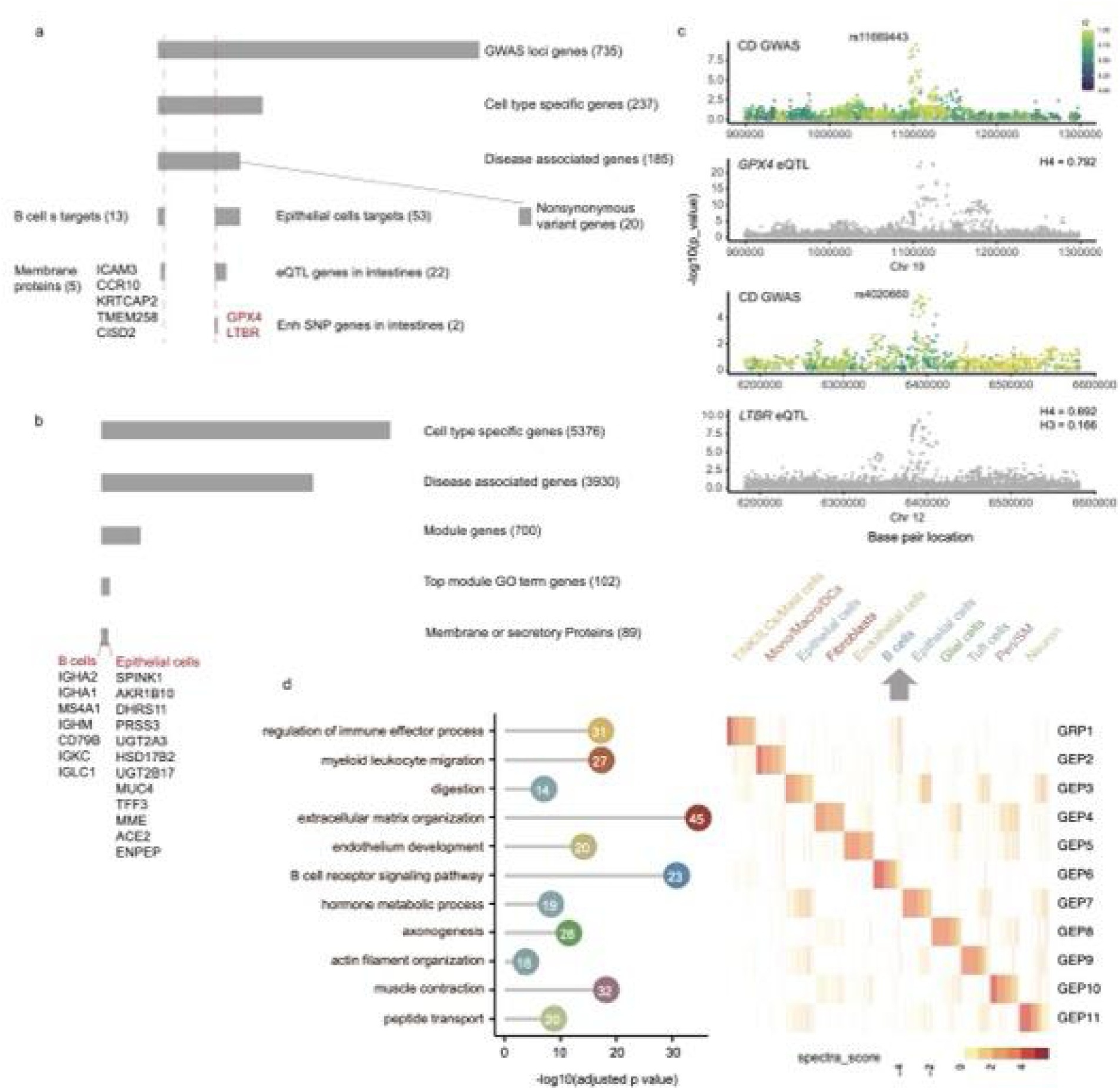
Identification of CD critical genes. **a-b**, Schematics of CD critical genes identification frameworks. **c**, Scatter plots showing genetic colocalizations between CD GWAS profiles and *GPX4*/ *LTBR* eQTL profiles. **d**, Left: Lollipop plot showing top enriched pathway in each gene expression program (GEP). Right: Heatmap showing spectra scores of top 200 GEP genes.

In the second approach, we first applied non-negative matrix factorization (NMF) to identify the core gene programs in CD inflamed tissues and matched each program to their involved cell types. In line with their cellular origins, cell subsets of the same cell lineages were clustered together (Extended Data Fig. 8d). For example, T cells and NK/ILC shared the same program associated with regulation of immune effector process and epithelial subsets enriched for gene programs related to digestion and hormone metabolic processes (Fig. 7d). Subsequently intersecting genes contributing to gene programs with cell type–specific genes and disease-associated genes, we acquired 700 genes with potential functional involvement in CD. Next, we kept 89 genes belonging to the top module pathway and coding membrane or secretory proteins (Fig. 7b, Supplementary Table 6). Notably, digestion related genes *MUC2*, *MUC4*, *SPINK1*, *TFF3* and *PRSS3*, and metabolism-related genes, *UGT2A3*, *UGT2B17*, *MME*, *AKR1B10*, *DHRS11*, *HSD17B2*, *ACE2* and *ENPEP* popped up as the most important functional genes for intestinal epithelial cells.

## Discussion

In this study, we established a high-quality single-cell atlas for human intestine tissues. With the data, we comprehensively investigated the differences between large and small intestines, and more importantly, cell states contributed to different clinical inflammatory phenotypes of IBD. Leveraging the single-cell transcriptional-level information, we illustrated the cellular and molecular mechanisms behind genetic regulation of CD risk and intestinal gene expression and provided rich computational evidence supporting the prioritization of CD critical cell types and genes, which may serve as treatment targets. For example, the pathological role of *GPX4* and *LTBR*, highlighted in our study, have been validated in mouse models. It was reported that an inflammatory response of Gpx4-deficient intestinal epithelial cells could be evoked by ω-6 polyunsaturated fatty acid^60^ and wild-type mice given LTBR inhibitors had an increased tumor burden in DSS induced colon cancer^61^. These examples support the value of our prioritization in IBD treatment targets identification.

The single-cell atlas contains >700,000 cells expressing >800 genes of 442 samples from 196 donors. Seventy-one cell subsets were annotated with distinct expression patterns. As a parallel study after CRC integrative atlas^16^, applying the same integrative and annotation strategies, our results presented a full blueprint of human intestinal immune-mediated diseases. It is interesting that highly contrasting results were observed for both cell occurrence and cell subset preference in CD and CRC, where the former represents a hyper-response context, while the latter represents an immune suppressive microenvironment. CRC was marked with enhanced cell occurrence in tumors compared to normal tissues^16^, whereas we reported reduced cell occurrence in CD. We speculated that cancer cells may force the other local cell types resulting in the observed phenotype by increasing cross cell type interactions. Therefore, it is surprising that a highly interactive yet not-sparse cell network was also found for CD. Together with the results that pericytes and fibroblasts served as the major senders in intercellular communication, we proposed that the acquired additional signaling from stromal cells caused dysregulation in CD.

Different from the stromal cell components that exhibited the largest alteration in CRC^16^, a strong cell functional subset remodeling was observed for immune cells in IBD, in line with that the immune cells being identified as the major target cells in the cell communication. We revealed the enrichment of Th1, *AXL*^+^*SIGLEC6*^+^ DCs (AS DCs), cDC1-derived LAMP3^+^ DCs and ABCs in IBD and more importantly, SPP1^+^ macrophages colocalized with FAP^+^ fibroblasts related to anti-TNF treatment resistance. The other explorations upon intra- and intro-cellular signaling provided references related to human intestinal immune regulations.

We took the advantage of integrative analysis in characterizing a rare but important cell type. Neuro-immune interactions have attracted extensive attention but limited by the sampling in human^6,62^. We have not only presented the proinflammatory states of neurons but also uncovered their expression patterns of major neuronal receptors in the human intestines, providing a window for understanding neuron-immune circuits in tissues.

The investigations of CD critical cell types and genes resulted from a systematic integration of multiple cell type specific transcriptomic and genetic profiles. Despite the fact that these pieces of evidence may point to different conclusions. We believe our efforts largely facilitated the comprehensive understanding of the cellular and molecular mechanisms under CD.

We must acknowledge possible confounders and limitations in this study. First, CD commonly occurs in early childhood, while the incidence of CRC increases along with aging. Thus, age might confound our comparisons between CD and CRC, which may see improvement in future studies. Second, sampling region (epithelial or lamina propria layers) could influence the cell composition of the samples. However, considering the commonly mixing of sampling, we evaluated their general characterization without controlling the sampling layers, which may introduce unwanted bias. Third, different from other cell subsets with exclusive expression patterns, the annotation of Tfr (n = 61 cells) was based on the co-expression of limited Tfh and Treg markers, leading to less confidence. Thus, we did not put any focus on this cell subset in this study and suggest a cautious interpretation for future applications. Lastly, minor effects such as genetic structural variation among different ancestries were not fully controlled, appealing for an in-depth investigation in future studies.

In conclusion, our study comprehensively characterized the single-cell landscape of IBD and proposed critical cell types and genes associated with CD pathology, providing insights in both cellular and molecular mechanisms of IBD and human intestinal immunity. The results may guide the development of preventive medicine, biomarkers, and new therapeutic targets for intestinal immune-mediated diseases.

## Code availability

This study did not generate any algorithm or software. Any additional information required to reanalyze the data reported in this paper is available from the lead contact upon request.

## Acknowledgments

We are grateful to Dr. Y. Miao and Dr. G. Liu for discussion. The presented study is supported by the Fundamental Research Funds for the Central Universities (Grant 10300310400209510) for X.C.

## Author contributions

X.C. designed the study. X.C. and S.W. collected the data and performed bioinformatic analysis supervised by E.P. and with the help from P.Y. J.W. performed IHC experiment. H.L. constructed the online browser with the support from B.Z. X.C. interpreted the data with the help from Q.Y. and Y.T. X.C. wrote the manuscript with input from all authors. The authors read and approved the final manuscript.

## Competing interests

The authors declare no competing interests.

## Notes

### Competing Interest Statement

The authors have declared no competing interest.

### Summary of Updates

Syntax errors have been corrected; author informations updated.

